# Asymmetric representation of symmetric semantic information in the human brain

**DOI:** 10.1101/2024.02.09.579613

**Authors:** Jiaxin Wang, Kiichi Kawahata, Antoine Blanc, Naoya Maeda, Shinji Nishimoto, Satoshi Nishida

## Abstract

Specific pairs of semantic entities have symmetric relationships, such as word pairs with opposite meanings (e.g., “intelligent” and “stupid”; “human” and “mechanical”). Such semantic symmetry is a key feature of semantic information. However, the representation of symmetric semantic information in the brain is not yet understood. Additionally, it is unclear whether symmetric pairs of semantic information do have symmetric representations in the brain? We addressed this question in a data-driven manner by using the voxelwise modeling of movie-evoked cortical response measured by functional magnetic resonance imaging. In this modeling, response in each voxel was predicted from semantic labels designated for each movie scene. The semantic labels consisted of 30 different items, including 15 pairs of semantically symmetric items. Each item was manually evaluated using a 5-point scale. By localizing the semantic representation associated with each item based on the voxelwise accuracy of brain-response predictions, we found that semantic representations of symmetric item pairs are broadly distributed but with little overlap in the cortex. Additionally, the weight of voxelwise models revealed highly complex, various patterns of cortical representations for each item pair. These results suggest that symmetric semantic information is rather asymmetric and heterogeneous representations in the human brain.

**Significance statement:** This study aimed to investigate if symmetric pairs of semantic information have symmetric representations in the human brain using statistical modeling of functional magnetic resonance imaging signals evoked by naturalistic movies. We built a model based on movie labeling of symmetric semantic items to quantify the cortical representations of symmetric semantic entities. These findings showed that symmetric pairs of semantic entities are represented in widespread cortical regions; however, still exhibit little overlap in localization and heterogeneous representations. These results offer significant insights into the cortical representations of semantic symmetry and advance the understanding of representational structure of semantic information in the human brain.

## Introduction

Semantic “symmetry” (or “oppositeness”) is a prominent feature of semantic information (Katz, 1972; Kempson, 1977). The symmetric semantic information is represented by word pairs with opposite meanings (i.e., antonyms; e.g., “masculine” and “feminine”) and effectively manipulated for high-level semantic processing by humans, such as thinking and communication. Regarding the neural mechanisms underlying symmetric semantic information processing, despite previous research on its generation and comprehension (Jeon et al., 2009; van de Weijer et al., 2014), its representation in the brain is not yet been fully understood. Therefore, there is no consensus as yet on whether or not symmetric information is represented symmetrically in the brain.

For comprehensively exploring semantic representations distributed across the brain, data-driven approaches based on voxelwise modeling of functional magnetic resonance imaging (fMRI) responses to naturalistic stimuli (e.g., movies and stories) have been widely employed (Huth et al., 2012, 2016; Güçlü and van Gerven, 2015; Vodrahalli et al., 2018; Deniz et al., 2019; Nishida et al., 2020a, 2021; Popham et al., 2021; Matsumoto et al., 2022). In this modeling framework, word features describing semantic information with naturalistic stimuli play a key role in determining the type of semantic information that can be modeled in the brain. Huth et al., who established such a data-driven approach for studying semantic representations, assigned noun/verb binary labels to each scene of movie stimuli as a simple word feature (Huth et al., 2012). This word feature is suitable for modeling cortical representations of concrete objects and actions but not for modeling semantic entities with abstract meaning or directionality, including symmetric semantic information.

More recent studies on voxelwise modeling have used word embeddings as an extension of binary labeling to efficiently model the structure of semantic representations in the brain (Güçlü and van Gerven, 2015; Huth et al., 2016; Vodrahalli et al., 2018; Nishida et al., 2020a, 2021; Popham et al., 2021; Matsumoto et al., 2022). Word embeddings are learned from large-scale text data using natural language processing algorithms, such as word2vec (Mikolov et al., 2013a), based on word co-occurrence statistics in sentences (Turney and Pantel, 2010). These word features are useful for modeling similarity structure of semantic representations (Mikolov et al., 2013b; Pereira et al., 2016; Nishida et al., 2021) but not for modeling their symmetric structure (Tang et al., 2014); note that “similarity” is used in a narrow sense here; although some researchers classify symmetry as sort of similarity (Turney and Pantel, 2010). Therefore, within the existing literature, the representation of symmetric semantic information in the brain has not yet been explored.

To investigate the representational characteristics of symmetric semantic information in the brain using a data-driven approach, we used the rating of symmetric word items as a word feature for voxelwise modeling of movie-evoked fMRI responses (Figure 1). In this rating, each scene of naturalistic movies used as experimental stimuli was manually evaluated on a 5-point scale for 30 different word items, including 15 pairs of semantically symmetric words (Table 1). We trained voxelwise models for predicting movie-evoked voxel response from rating of each item. Subsequently, the model performance of response prediction for each item was tested to identify localization of cortical semantic representations associated with each symmetric item pair. Finally, we visualized the cortical distribution of model weights for each item to assess representational patterns of each symmetric item pair.

**Figure 1.**
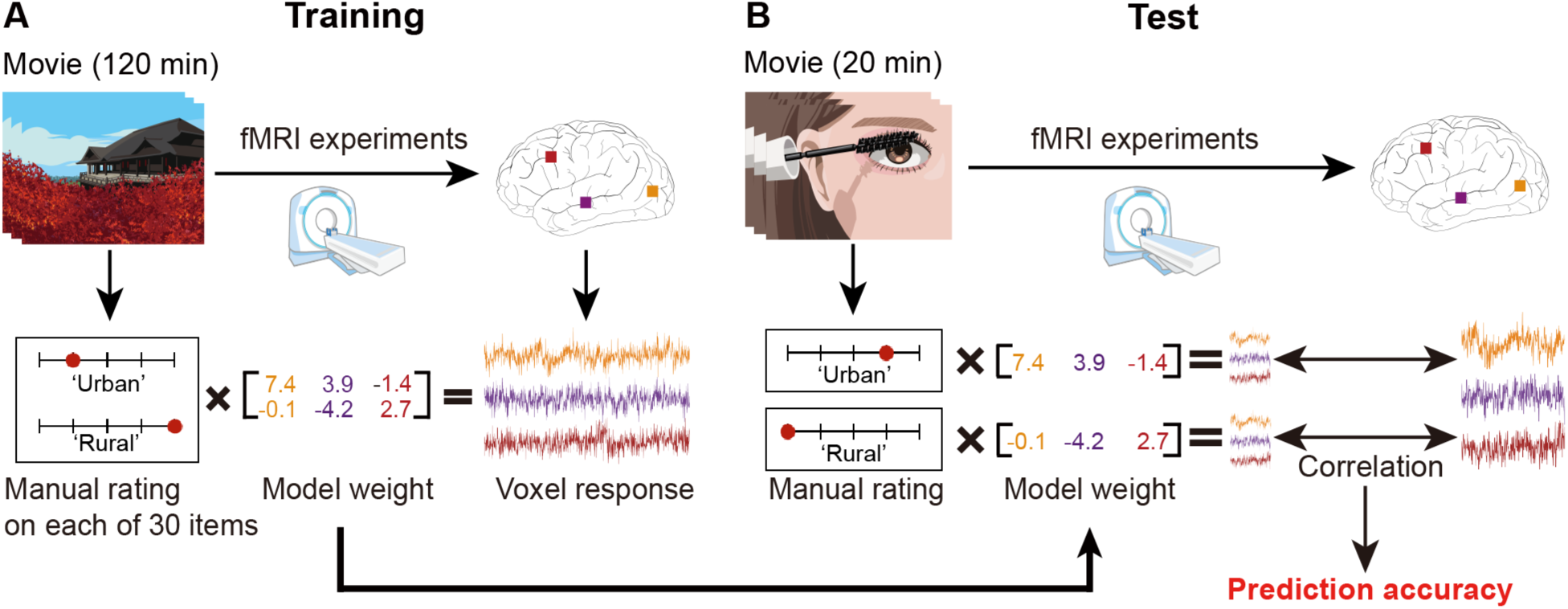
Voxelwise modeling. **A**, Model training. To train models for brain-response prediction, we collected voxel responses to natural movie scenes and the manual rating of the scene on each of 30 different word items, including 15 symmetric pairs (Table 1). A linear regression model was trained to predict the response from each participant in each voxel from the manual rating of the scenes. Each value in the model weights reflects bidirectional selectivity for each item within each voxel. **B**, Model test. The prediction accuracy in each voxel was evaluated using the Pearson correlation coefficient between predicted and measured voxel responses to new movie scenes. To compute the prediction accuracy for each of the 30 items, the model weights were decomposed into those for each item and used to predict the voxel response from the manual rating of the item.

**Table 1.** Rated word items and their symmetric pairs used for the analysis.

| Item | Paired item |
| --- | --- |
| Beautiful | Ugly |
| Urban | Rural |
| Nauseating | Cute |
| Cheap | Lush |
| Masculine | Feminine |
| Amusing | Gloomy |
| Modern | Traditional |
| Simple | Complex |
| Warm | Cool |
| Sensitive | Bold |
| Intelligent | Stupid |
| Static | Dynamic |
| Human | Mechanical |
| Clean | Filthy |
| Noisy | Quiet |

## Materials and Methods

### Participants

A total of 52 healthy Japanese participants (21 females and 31 males; aged 20–61 years, mean ± SD = 26.8 ± 8.9 years) were recruited for two sets of fMRI experiments. Among them, 36 participated only in one of the fMRI experiments, and 16 participated in both. All participants had normal or corrected-to-normal vision. Written informed consent was obtained from all participants. The experimental protocol was approved by the ethics and safety committees of the National Institute of Information and Communications Technology.

### Experimental design

In each fMRI experiment, the participants were asked to view Japanese web or TV ad movies with sound on a projector screen inside the scanner. The web and TV ads movie sets included 368 and 420 ads, respectively, which were all unique and consisted of a wide variety of product categories (Nishida et al., 2020b). The length of each movie was typically 15 or 30 s. To create the movie stimulus for each experiment, the original movies in each movie set were sequentially concatenated in a pseudorandom order. For each movie set, we obtained 14 nonoverlapping movie clips with duration of 610 s.

Individual movie clips were then displayed in separate fMRI runs. The initial 10 s of each clip served as a dummy to discard hemodynamic transient signals caused by clip onset. The voxel responses collected during this dummy phase were not used for modeling. Twelve clips from each movie set were presented only once and voxel responses to these clips were used for training voxelwise models (training dataset; 7200 s in total). The other two clips for each movie set were presented four times each in four separate fMRI runs and voxel responses to these clips were averaged across runs to improve the signal-to-noise ratio (Nishimoto et al., 2011; Huth et al., 2012; Nishida and Nishimoto, 2018). The averaged responses were further used to test voxelwise models (test dataset; 1200 s).

### MRI data collection and preprocessing

The MRI data and preprocessing techniques used in this study are the same as previously reported for different research objectives (Nishida et al., 2020b, 2021; Kawahata et al., 2022) and are summarized in this section. fMRI data were collected through a 3T Siemens MAGNETOM Prisma scanner (Siemens, Germany) with a 64-channel Siemens volume coil, using a multiband gradient echo EPI sequence (Moeller et al., 2010; TR = 1000 ms; TE = 30 ms; flip angle = 60°; voxel size = 2 × 2 × 2 mm; matrix size = 96 × 96; number of slices = 72; multiband factor = 6). Additionally, anatomical data were gathered using a T1-weighted MPRAGE sequence (TR = 2530 ms; TE = 3.26 ms; flip angle = 9°; voxel size = 1 × 1 × 1 mm; matrix size = 256 × 256; number of slices = 208) on the same 3T scanner. The fMRI data for each participant were collected in three separate recording sessions over three days for each of the two sets of fMRI experiments.

Motion correction in each fMRI run was conducted using the statistical parameter mapping toolbox (SPM8, http://www.fil.ion.ucl.ac.uk/spm/software/ spm8/). All volumes were aligned to the first image from the first run for each subject. The low-frequency voxel response drift was then eliminated by subtracting median-filtered signals (within a 120-s window) from raw signals. The response for each voxel was normalized by subtracting the mean response and scaling to the unit variance. FreeSurfer (Dale et al., 1999; Fischl et al., 1999) was used for identifying cortical surfaces from anatomical data and registering these surfaces to the voxels of functional data. Subsequently, each voxel was assigned to one of the 356 cortical regions derived from the HCP-MMP1 atlas for cortical segmentation (Glasser et al., 2016).

### Statistical analyses

#### Manual rating of movie scenes

We constructed a feature space of symmetric semantic information from the manual rating of movie scenes on 30 different word items consisting of 15 pairs with opposite meanings (Table 1). Multiple raters manually rated every 2-s scene of each movie set. The raters for web ad movies completely differed from the fMRI participants (26 females and 6 males; aged 24–61 years). Conversely, the raters for TV ad movies partly overlapped with the fMRI participants (3 females and 3 males [aged 22–46 years] among fMRI participants; 5 females and 1 male [aged 20–45 years] from others). Although the raters sequentially watched separate 2-s scenes of the movies on a computer screen, they rated each item on a 5-point scale from 0 to 4. To maintain the raters’ motivation, only 15 items were assigned to each in a single-scene rating. To ensure cross-subjectivity of the rating, each scene in web ad movies was assessed by nine different raters, whereas in TV ad movies each scene was evaluated by 12 different raters. Next, the ratings for each 2-s scene were averaged separately for each movie set. Finally, the average ratings for each item were oversampled to obtain a time series of ratings in every 1-s scene that matched the sampling rate of fMRI data.

#### Voxelwise modeling

We performed voxelwise modeling (Naselaris et al., 2011) using the symmetric semantic information feature space. A series of voxel responses to movie scenes in each participant were modeled as a weighted linear combination of the features (i.e., average ratings of movie scenes). To capture the slow hemodynamic response and its coupling with voxel response (Nishimoto et al., 2011), four sets of features with delays of 3, 4, 5, and 6 s were concatenated, yielding 120 features (30 items × 4 delays). The linear weights of the model were estimated using an L2-regularized linear least-squares regression, which can obtain good estimates in voxelwise modeling (Huth et al., 2012).

We estimated the regularization parameter of each model by randomly dividing the training dataset into two subsets, comprising 80% and 20% of samples for model fitting and validation, respectively. This random resampling procedure was repeated 10 times. The regularization parameters were optimized based on the mean Pearson correlation coefficient between the predicted and measured voxel responses for the 20% validation samples.

#### Evaluation of model performance

For examining the predictability of movie-evoked brain responses by the trained voxelwise models, the performance of model in terms of voxel-response prediction (prediction accuracy) was evaluated using the test dataset that was not used for model training. The prediction accuracy in each voxel was quantified as the Pearson correlation coefficient between the predicted and measured responses in that voxel. The prediction accuracy of each model was computed using all the 30 word items (whole-item prediction accuracy) as well as each item (item-wise prediction accuracy). The item-wise prediction accuracy was quantified using the Pearson correlation coefficient between the predicted and measured voxel responses when the voxel response was predicted from a single item and the corresponding model weights. The P-value associated with the prediction accuracy in each voxel was corrected for multiple comparisons using false discovery rate (FDR).

To explore the cortical distribution of symmetric-pair representations and their overlap, we evaluated the localization of cortical voxels exhibiting significant item-wise prediction accuracy for each item pair per participant (P < 0.05, FDR corrected). For each item pair, we calculated the fraction of voxels in which item-wise prediction accuracy was significant for either one or both items, after excluding voxels in which prediction accuracy was not significant for either of the items. Specifically, when considering item pairs A and B, we counted the number of voxels with significant prediction accuracy for both A and B, denoted as *N*A∩B and for any of A and B, denoted as *N*A∪B. The fraction of overlapping voxels was then calculated by *N*A∩B/*N*A∪B.

The fraction of overlapping voxels for each item pair was compared with its corresponding chance level to assess if the observed voxel overlap significantly deviates from chance. The chance level was computed as follows: for a given item pair A and B, all voxels with significant prediction accuracy for any of A and B, denoted as *V*A∪B, were extracted. Next, the assignment of voxels with significant accuracy for only B was randomly shuffled among *V*A∪B, and the overlapping voxels fraction was calculated. This procedure was repeated 1000 times and the average fraction of overlapping voxels over these repetitions provided the chance level for the fraction of overlapping voxels specific to the A and B pair.

For group-level analysis of model performance, we merged the voxelwise models of individual participants across movie sets, unless indicated otherwise. Nevertheless, highly similar trends were observed in results even when analyzing models separately for each movie set (see Results).

#### Evaluation of the model weights

The weights of the voxelwise model, which reflects linear selectivity of each voxel for semantic information associated with each word item, can be considered as cortical representation of such semantic information. To explore the heterogeneity/homogeneity of the cortical representation of symmetric semantic information, we evaluated the pattern of distribution of model weights across the cortex for each symmetric word pair.

Each voxel had four weight values for each item and captured hemodynamic response delays (see voxelwise modeling); the four values were averaged to obtain a single weight value for each item. Additionally, the voxel was tested for the significance of its prediction accuracy for the item (P < 0.05, FDR corrected). The item-wise representation in each voxel was classified into three types based on the combination of the weight sign and significance of prediction accuracy. A voxel displaying a positive (negative) weight and significant prediction accuracy was considered to have a positive (negative) representation for the item. Conversely, a voxel with non-significant prediction accuracy was considered to have no representation for the item.

Subsequently, the item-pair representation in each voxel was classified into eight types as per the combination of item-wise representations for each symmetric item pair: positive–positive, positive– no, positive–negative, no–negative, negative–negative, negative–no, negative–positive, and no– positive representation pairs. Finally, for each item pair, the fraction of voxels classified into each of these eight types was separately calculated for each participant after excluding voxels with non-significant prediction accuracy for both items.

For group-level analysis of model weights, individual models were merged across movie sets, unless indicated otherwise. Nevertheless, we observed highly similar trends of results when the movie sets were separately analyzed even in this case (see Results).

## Results

### Symmetry of ratings for each word pair

This study investigated the neural representation of symmetric semantic information associated with 15 word pairs (Table 1). Before examining neural representation, we aimed to confirm that these word pairs actually reflect semantic symmetry in human perception. If so, the manual ratings of movie scenes for individual word items would exhibit a negative correlation across each word pair. To test this hypothesis, we initially evaluated the Pearson correlation coefficient between the time series of manual ratings for each item and its paired item. Overall, we found significantly negative correlations for paired items in both movie sets (Figure 2; web ad movies: mean r = −0.26; TV ad movies: mean r = −0.39; one-sample t-test, P < 10^−5^). In contrast, we observed no significant or significantly positive correlations for unpaired items (web ad movies: mean r = 0.04; TV ad movies: mean r = 0.01; P <10^−4^ and 0.32 respectively). For each pair, there were significantly negative correlations (Figure 2-1; P < 10^−23^, FDR corrected) except for two pairs (“warm” and “cool,” and “sensitive” and “bold”). These results indicate that the selected word pairs capture the human perception of semantic symmetry.

**Figure 2.**
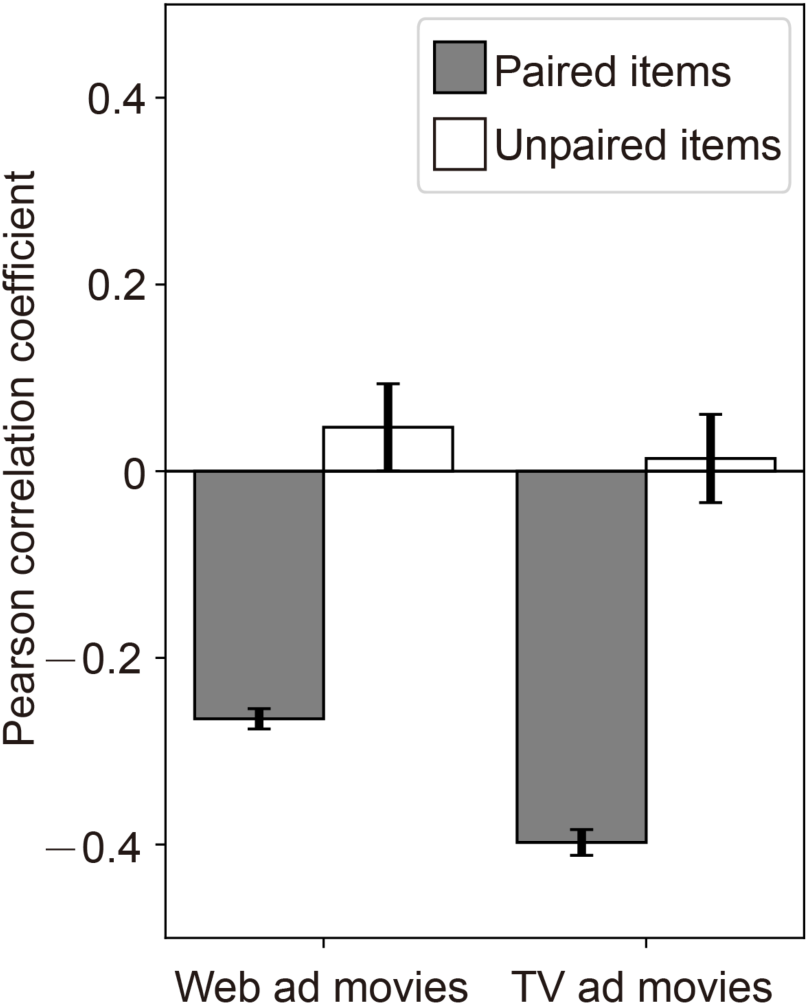
Negative correlations of ratings for symmetric item pairs. To examine the perceptual symmetry of word item pairs used for voxelwise modeling (Table 1), we computed the Pearson correlation between a time series of manual ratings for each item and its paired item. In addition, as baseline, we calculated the correlation for all possible combinations of unpaired items. Gray and white bars represent the Pearson correlation coefficients averaged over all paired and unpaired items, respectively, separately shown for the web and TV ad movie sets (error bars, standard error of the mean [SEM]). For the correlation between each symmetric item pair, see Figure 2-1.

### Voxelwise models predict movie-evoked responses in widespread cortical regions

Previous studies have demonstrated that voxelwise models with semantic features associated with thousands of words broadly predict stimulus-evoked responses across the cortex (Huth et al., 2012, 2016; Nishida et al., 2021; Popham et al., 2021). To examine whether our semantic voxelwise model based on the rating of only 30 word items exhibit a similar widespread distribution of predictable voxels across the cortex, we calculated the whole-item prediction accuracy of the models (i.e., when using all 30 items) similarly to previous studies (see Materials and Methods).

Predictable voxels were distributed in widespread cortical regions not only in a single participant (Figure 3A) but in the entire population (Figure 3B). High prediction accuracy was observed mainly in occipitotemporal and superior temporal cortical regions, which include object-selective regions (Malach et al., 1995; Kanwisher et al., 1997; Epstein and Kanwisher, 1998; Downing et al., 2001), which is consistent with previous reports on semantic voxelwise modeling of movie-evoked responses (Huth et al., 2012; Nishida et al., 2021). This tendency was preserved when each movie set was separately analyzed (Figure 3-1). Thus, our voxelwise models based on only 30 word items can predict movie-evoked responses broadly across a wide range of the cortex.

**Figure 3.**
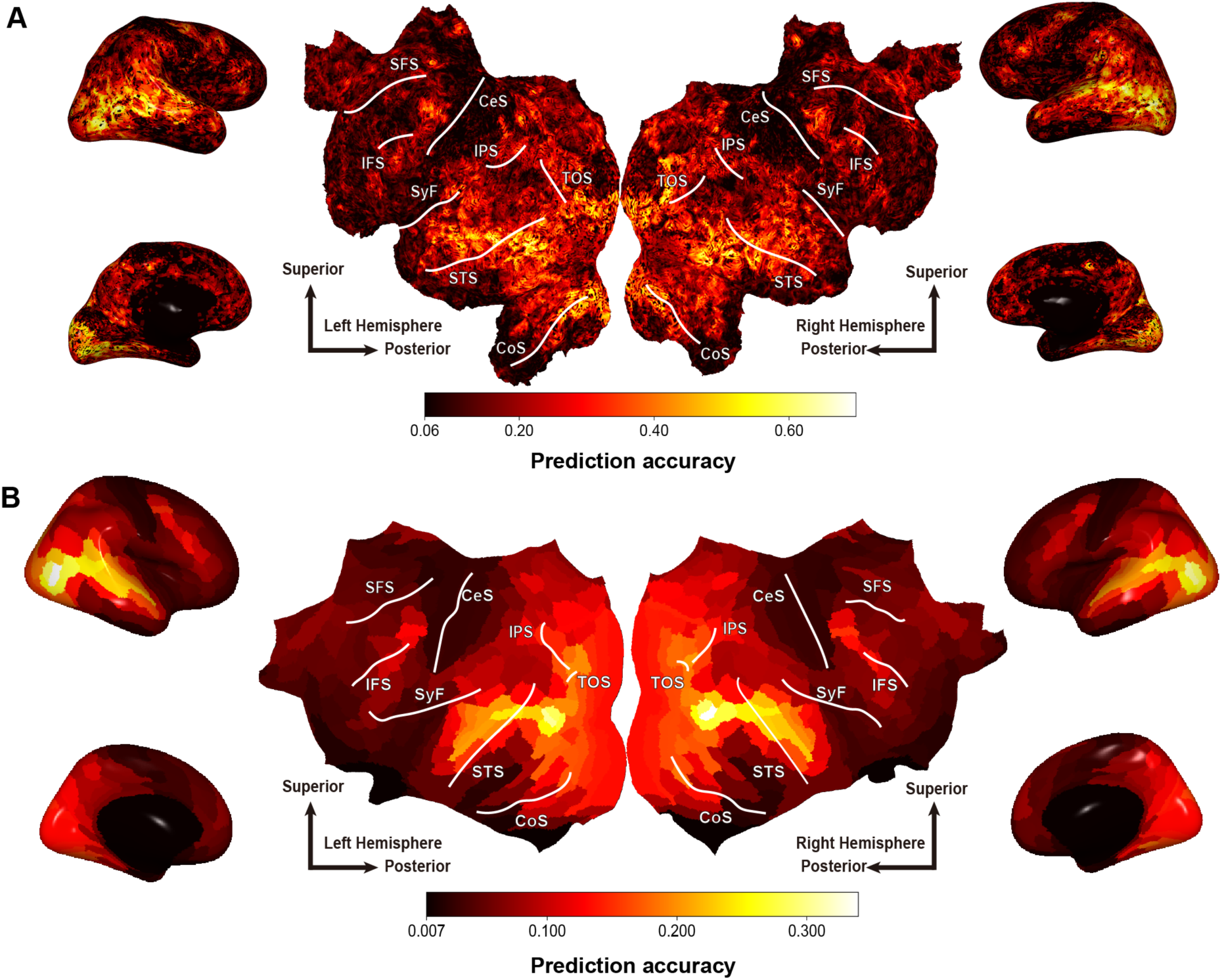
Cortical mapping of the prediction accuracy in voxelwise models. **A**, Whole-item prediction accuracy in each voxel for a representative participant, mapped onto their cortical surface. This map was derived from the web movie set. Brighter colors on the map indicate voxels with higher prediction accuracy. Only voxels with prediction accuracy over a significance threshold (P < 0.05, FDR corrected) are shown. White lines on the cortical maps denote representative sulci: CoS, collateral sulcus; STS, superior temporal sulcus; TOS, transverse occipital sulcus; IPS, intraparietal sulcus; SyF, sylvian fissure; CeS, central sulcus; SFS, superior frontal sulcus; IFS, inferior frontal sulcus. **B**, Region-wise whole-item prediction accuracy for both movie sets mapped onto the cortical surface of a reference brain. After averaging the voxelwise prediction accuracy of each participant’s model within each cortical region, the region-wise whole-item prediction accuracy was averaged across participants. Brighter colors on the map indicate regions with higher prediction accuracy. Only regions with prediction accuracy over a significance threshold (P < 0.05, FDR corrected) are shown. All other conventions are as in **A**. For the prediction accuracy in each movie set, see Figure 3-1. For item-wise prediction accuracy, see Figures 3-2 to 3-4.

We also quantified item-wise prediction accuracy, i.e., model performance of voxel-response prediction for each word item. Despite the varying prediction accuracy and cortical distribution from item to item (Figures 3-2 to 3-4), the fraction of significantly predictable voxels in the whole cortex was sufficiently high (Figure 4; 19.7–46.4% [mean = 34.0%] of all cortical voxels). This indicates that our voxelwise models based on scene ratings for word items can effectively capture the semantic representations associated with those word items.

**Figure 4.**
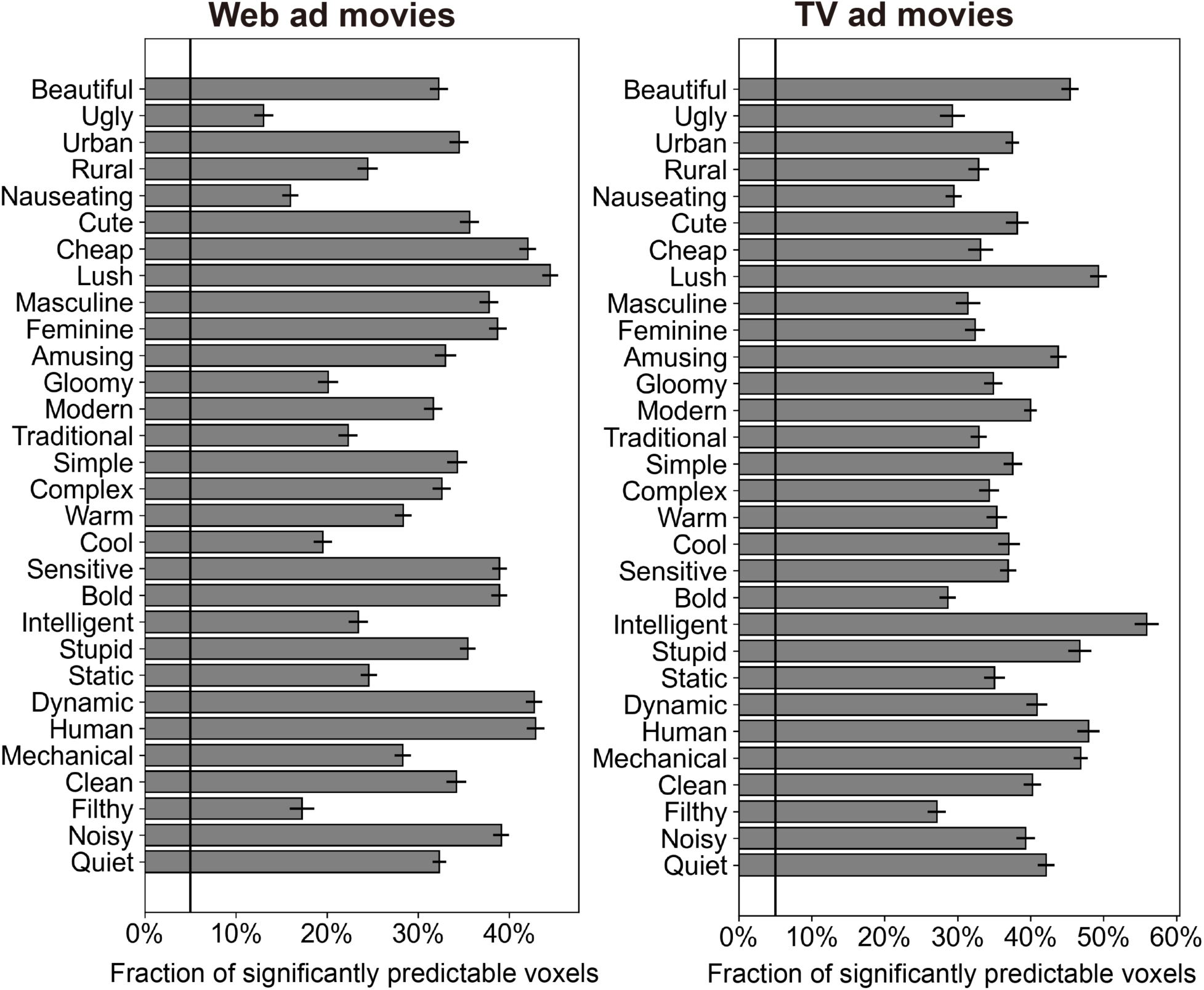
Fraction of significantly predictable voxels in the entire cortex for the 30 word items. We tallied the number of voxels exhibiting significant prediction accuracy (P < 0.05, FDR corrected) for every word item and each participant. We then calculated the fraction of significantly predictable voxels by dividing this count by the total number of cortical voxels. The mean fraction of significantly predictable voxels across participants for each item is shown separately for each movie set (left, web ad movies; right, TV ad movies; bars, mean fraction; error bars, SEM). All these fractions were significantly higher than the significance threshold of 0.05; denoted by the horizontal dashed line; one-sample t-test, P < 10^−10^, FDR corrected).

### Cortical representations of semantically symmetric pairs rarely overlap

If the symmetric pairs of semantic information were represented symmetrically in specific regions of the cortex, the voxels predicted by our voxelwise model should largely overlap within each symmetric item pair. To examine this possibility, we calculated the overlap fraction of voxels with significant prediction accuracy (P < 0.05, FDR corrected) for each item pair.

However, we rarely observed such overlapping voxels (Figure 5). In a representative participant, although each item in an example symmetric pair, “human” and “mechanical,” exhibited predictable voxel distributions across widespread cortical regions, there was little overlap among their predictable voxels (Figure 5A). At the group level, there was only 18.2%–46.6% (mean = 27.9%) overlap among significantly predictable voxels per item pair (Figure 4B). A similar tendency was observed for each movie set (Figure 5-1).

**Figure 5.**
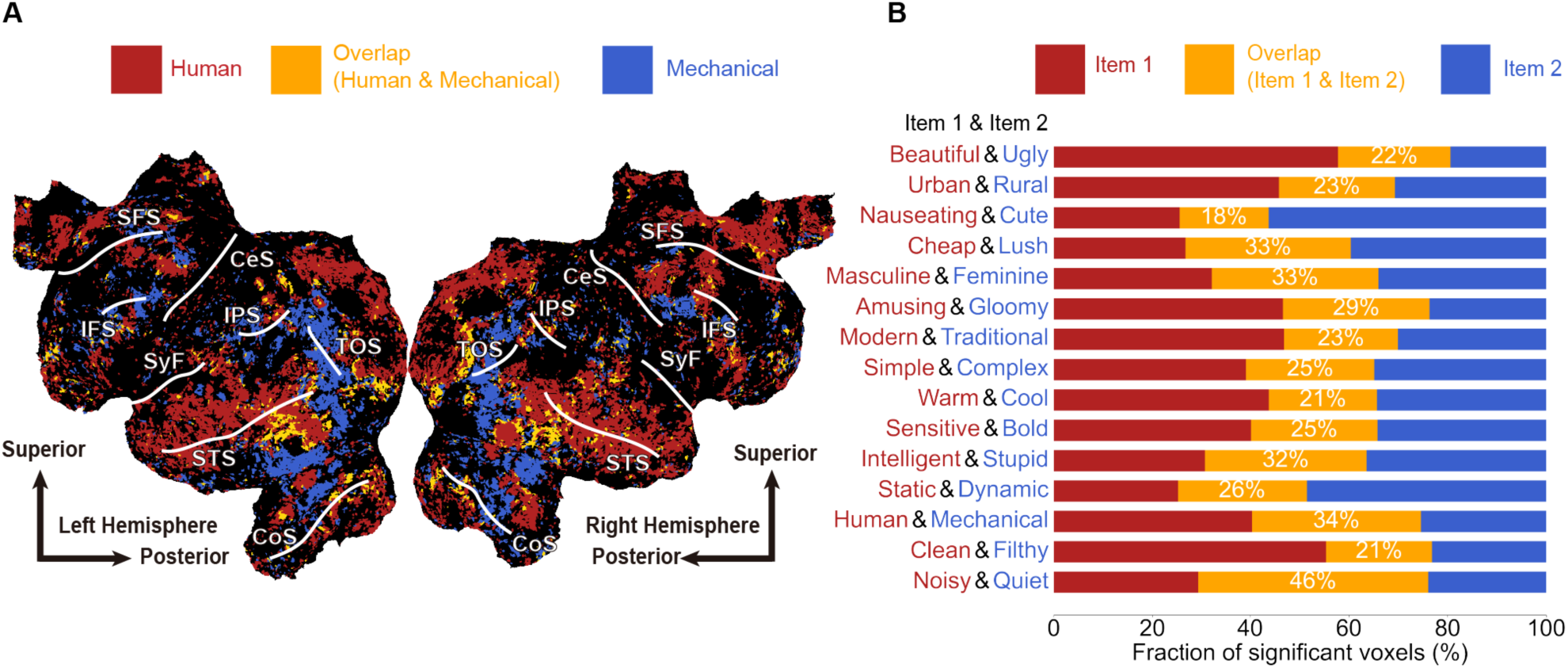
Little overlap of predictable voxels for semantically symmetric pairs. **A**, Example item pair exhibiting little overlap among their predictable voxels. This result was derived from a representative participant in the web movie set. Voxels in which “human” (red) and/or “mechanical” (blue) showed significant prediction accuracy (P < 0.05, FDR corrected) were mapped onto the participant’s cortical surface. Although the predictable voxels for each of these two items were broadly distributed across the cortex, overlap (orange) was rarely observed (24% of predictable voxels). Other conventions are the same as in Figure 3A. **B**, Fraction of different-type predictable voxels averaged over participants in both movie sets. For each item pair (items 1 and 2), the fraction of voxels predicted by either item 1 (red) or 2 (blue) and voxels predicted by both (orange) were separately calculated and averaged over all participants. The percentage denoted in each bar indicates the fraction of voxels predicted by both items (i.e., overlapping voxels). For the fraction in each movie set, see Figure 5-1. For control analysis with consideration of a collinearity problem, see Figure 5-2.

By comparing the observed fraction of overlap with its chance level (see Materials and Methods for how to calculate the chance level), we found that the observed fraction of overlap for all symmetric pairs was significantly lower than chance (Figure 6A; paired t-test, P < 10^−38^, FDR corrected). This trend remained when the comparison was performed for each item pair for individual participants (Figure 6-1), suggesting that the infrequent representational overlap of symmetric information is a common characteristic across item pairs.

**Figure 6.**
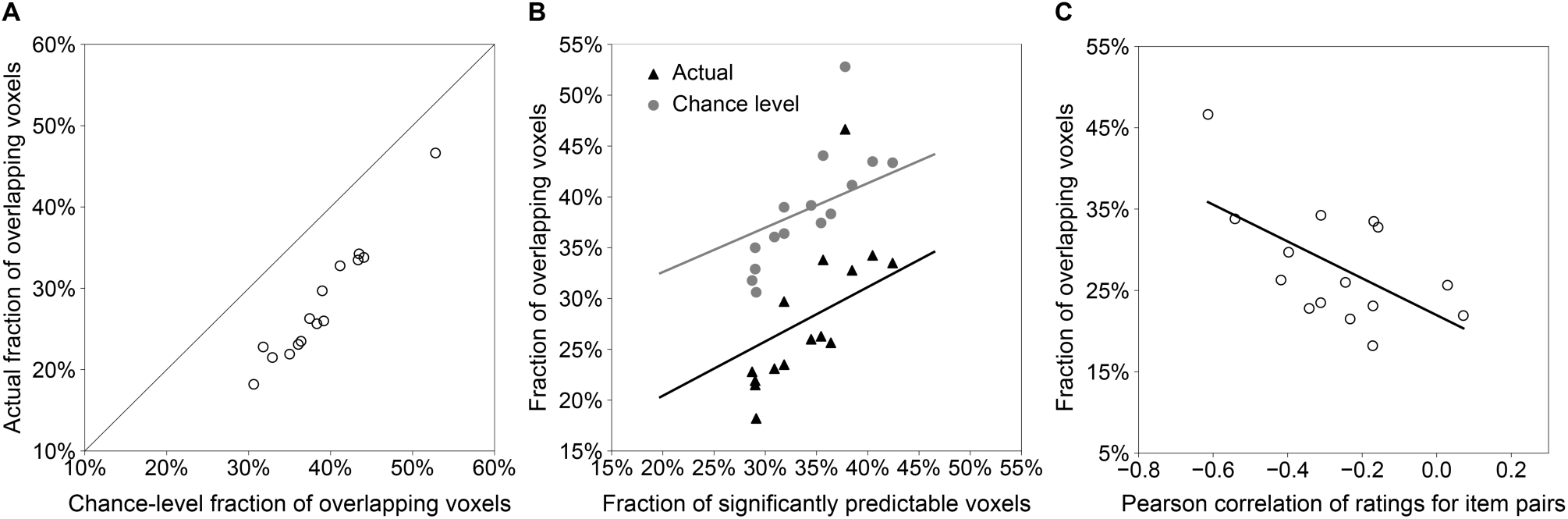
Fraction of overlapping voxels with respect to chance level, model prediction performance, and perceptual symmetry. **A**, Observed and chance-level fractions of overlapping voxels. The fraction of overlapping voxels, averaged over participants in both movie sets, is shown for each item pair compared with its chance level. Each circle represents the observed (y-axis) and corresponding chance-level (x-axis) fractions for an individual item pair. **B**, Correlation between fractions of overlapping voxels and the model prediction performance. The observed (black triangles) and chance-level (gray circles) fractions of overlapping voxels (y-axis) calculated in **A**, are plotted against the fraction of significantly predictable voxels (x-axis) for the corresponding pair. Black and gray lines indicate linear fits to data for the observed and chance-level fractions, respectively. **C**, Correlation between the fractions of overlapping voxels and perceptual symmetry. The observed fraction of overlapping voxels, calculated in **A**, is plotted against the Pearson correlation of manual ratings for each item pair, as in Figures 2 and 2-1. Each circle represents a single item pair. The line indicates a linear fit to the data.

The observed and chance-level fraction of overlap among item pairs (Figure 6A) appeared to be linked to the voxel-response prediction performance for each pair (Figure 6B). Both fractions were significantly correlated with prediction performance (Pearson r = 0.74 and 0.77, P < 0.01 and 0.001 for the observed and chance-level fractions, respectively). However, the fraction of overlap below the chance level persisted for the item pair with the highest prediction performance (“human” and “mechanical”). Further, the rate of change in these fractions in relation to prediction performance was similar for both observed and chance-level fractions (slope of linear fits = 0.53 and 0.43 for the observed and chance-level fractions, respectively). Consequently, the infrequent overlap observed is unlikely to stem from the poor performance of voxelwise models.

Nevertheless, further analysis revealed a behavioral correlate: the fraction of overlap was relatively higher for item pairs with stronger perceptual symmetry. We observed a significant correlation between the observed fraction of overlapping voxels and the manual ratings for each item pair (Figure 6C; Pearson r = −0.58, P < 0.05). This result indicates that despite the generally infrequent overlap across item pairs, those with stronger perceptual symmetry are more likely represented concurrently in the same voxels.

In our voxelwise modeling, we modeled voxel responses by linear combination of rating for multiple word items, including semantically symmetric pairs. The rating scores of these symmetric pairs showed negative correlations (Figure 2). This collinearity of regressors might lead to the rare overlap of predictable voxels for symmetric items, although the L2 regularization employed in our modeling effectively avoids the collinearity problem (Gruber, 2017). To test this possibility, we constructed voxelwise models separately using the rating of each item and calculated the overlap fraction of predictable voxels for each symmetric pair. Consequently, 30 different models, corresponding to 30 items, were obtained for each participant and movie set. Under these conditions, the overlap fraction was below the chance level (Figure 5-2), indicating that the collinearity of regressors has little impact on the rare overlap of predictable cortical voxels observed in our study.

### Cortical representations of semantically symmetric pairs are highly heterogeneous

The weights of the voxelwise models correspond to the linear selectivity of each voxel for the semantic information linked with each word item. This selectivity can be regarded as the representation of semantic information in that voxel. Therefore, to explore the cortical representations of semantically symmetric information, we visualized the distributional patterns of model weights across the cortex. If symmetric semantic information was symmetrically represented in the cortex, the model weights would be homogeneously distributed. Conversely, if represented asymmetrically, the model weights would show heterogeneous distributional patterns.

Our results showed highly heterogeneous patterns of model weights across the cortex (Figure 7). Next, the model weight of each voxel was classified into eight types according to its positive/negative/no selectivity for each symmetric item pair (see Materials and Methods). For the symmetric pair of “human” and “mechanical,” the eight types of model weights were intermingled across the cortex of a representative participant (Figure 7A). The group-average fraction of voxels classified into these eight types similarly revealed intermingled patterns of model weights for symmetric paired items in the cortex (Figure 7B). A similar tendency was observed for each movie set (Figure 7-1).

**Figure 7.**
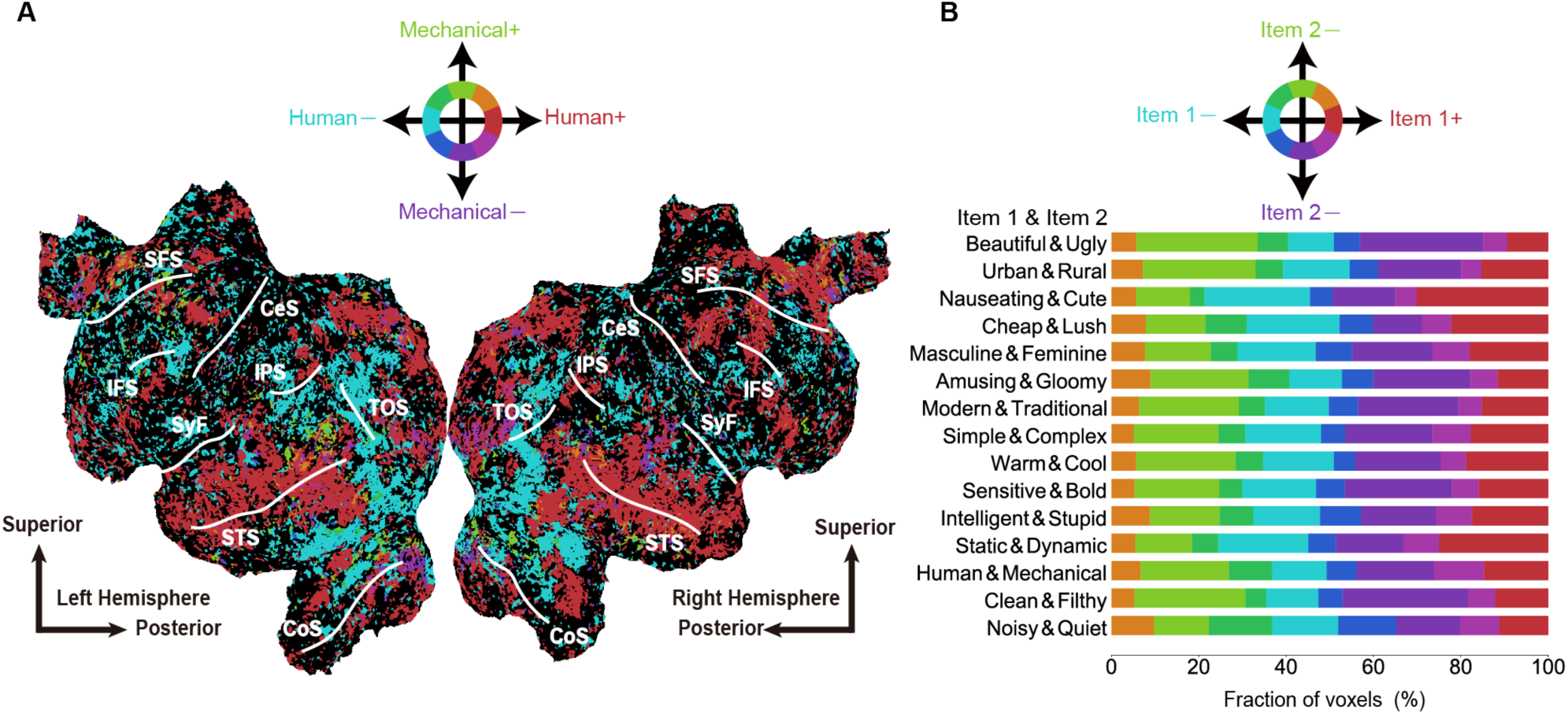
Heterogeneous cortical distribution of voxel selectivity for semantically symmetric pairs. **A**, Example cortical distribution of model weights in a representative participant from the web ad movie set. The model weight at each voxel was classified into eight types according to its positive/negative/no selectivity and mapped onto the participant’s cortical surface. The eight-type model weights are indicated using the HSV color code shown at the top. Other conventions were as those in Figure 3A. **B**, Fraction of different-type selective voxels averaged over participants from both movie sets. Bars indicate the voxel fraction of each type of selectivity for each symmetric item pair. The same HSV color code is used as in A. For each movie set, see Figure 7-1. For the distribution of fractions among participants, see Figures 7–2.

There are two potential explanations for the observed heterogeneous patterns of model weights at the group level. The more straightforward explanation is that individual brains have similar heterogeneous patterns. Another is that the pattern of model weights is homogeneous in each brain but varies across brains. However, the distribution of the model weight fraction for individual participants showed a similar fraction of each type of model weight across individuals (Figure 7-2). Thus, the heterogeneous cortical distribution of model weights appears to be a common characteristic even at the individual level.

## Discussion

To investigate how symmetric semantic information is represented in human brain, we quantified the cortical representations of such information using voxelwise modeling based on the movie-scene rating for 15 semantically symmetric pairs of items. Our voxelwise model predicted movie-evoked voxel responses in a sufficiently large portion of the cortex (Figures 3 and 4). Despite the broad distribution of predictable voxels were linked with each item across the cortex, there was little overlap of predictable voxels between each symmetric pair of items (Figures 5 and 6). The distribution of voxelwise model weights revealed highly heterogeneous representations of symmetric semantic information across the cortex (Figure 7). These findings suggest that symmetric semantic information is represented in the human brain in a distributed and asymmetric manner.

Voxelwise modeling of fMRI responses to naturalistic stimuli based on word features has been extensively used to investigate cortical distributed representations of semantic information associated with thousands of words for data-driven approaches (Huth et al., 2012; Güçlü and van Gerven, 2015; Huth et al., 2016; Vodrahalli et al., 2018; Deniz et al., 2019; Nishida et al., 2020a, 2021; Popham et al., 2021; Matsumoto et al., 2022). Word binary labeling to naturalistic stimuli is effective for modeling the discrete and categorical representations of objects and actions (Huth et al., 2012) and word embedding, which captures the rich semantic content of stimuli, is effective for modeling the similarity structures of semantic representations (Güçlü and van Gerven, 2015; Huth et al., 2016; Vodrahalli et al., 2018; Nishida et al., 2020a, 2021; Popham et al., 2021; Matsumoto et al., 2022). Although these word features are valuable for investigating the essential properties of cortical semantic representations, the stimulus ratings for symmetric word items used in this study are relatively more appropriate for modeling the representational structures of symmetric semantic information, which are important properties of semantic representations. To the best of our knowledge, this is the first study that explicitly explores the cortical representations of symmetric semantic information distributed across the cortex in a data-driven manner.

Our results demonstrate that the cortical representation of semantic entities shows little overlap between symmetric pairs at the voxel level (Figures 5 and 6). Thus, the representation of semantic symmetry is globally represented by interregional networks in the brain. This perspective is coherent with the widely accepted notion that semantic information is broadly distributed throughout the cortex (Martin, 2007; Patterson et al., 2007), as previously reported by voxelwise modeling of semantic representations (Huth et al., 2012, 2016). This study advances our understanding of cortical representations of semantic information by showing that even symmetric semantic entities possess distributed representations in the human cortex.

The cases in which symmetrical information is represented through interregional networks in the brain are not exclusively limited to semantic information. For instance, previous studies on the neural basis of emotion reported that positive and negative emotions are portrayed in different brain areas. Specifically, positive emotions (e.g., happiness and joy) are predominantly associated with the medial prefrontal cortex (Aharon et al., 2001; Ishai et al., 2004), whereas negative emotions (e.g., fear and disgust) are primarily linked with the amygdala and insular cortex (Phelps et al., 2001; Wicker et al., 2003). Therefore, this interregional representation of symmetrical concepts may be a common characteristic among some modalities of information representations in the human brain.

The use of a nonoverlapping interregional representation of symmetric information provides several advantages. One of the advantages is its flexibility in treating symmetry depending on the cognitive context. For instance, it may be more effective to treat symmetric pairs, such as beautiful and ugly, as opposing concepts and to accentuate their symmetry through regional interaction. However, such treatment may not be appropriate in some situations. Consider Francis Bacon’s paintings, often considered both ugly and beautiful. In such cases, reducing the effect of symmetry and allowing the co-occurrence of each representation of the pair may be essential. Modulating the effect of symmetry can be achieved by regulating interregional connections (van den Heuvel and Sporns, 2013; Shine et al., 2019), which is more effective than having symmetrical representations in a specific region. Therefore, an interregional representation of symmetric information offers a flexible approach to the treatment of symmetry based on contextual requirements. Modulating the impact of symmetry through interregional regulation allows for a nuanced representation of the underlying concepts.

This study has some limitations. One is related to the use of a linear regression model for voxelwise modeling. Although this approach has been effective in identifying those cortical semantic representations linearly associated with ratings of symmetric word items, it may not perform as well when the association is nonlinear. Nevertheless, using nonlinear models for voxelwise modeling is not advisable because of their lack of interpretability (Naselaris et al., 2011). Further, nonlinear models cannot easily provide interpretable information about model weights, as illustrated in Figure 7. However, our results do not preclude the possibility of a nonlinear representation of semantic symmetry in the cortex, and this will be the focus of our future investigations.

Another limitation of this study is that the cortical representations of semantic symmetry identified at the voxel level may not completely capture potential overlap and homogeneity at the level of individual neurons or local neural circuits. This raises the possibility that such representations exist at a more fine-grained scale but are not reflected in the macroscopic (voxel-level) signals in our analysis. However, it is widely accepted that voxel-level signals remain informative, as any bias in the microscopic representation tends to be expressed in the voxel signal (Boynton, 2005; Haynes and Rees, 2005; Kamitani and Tong, 2005). In other words, regions with dominant neurons and neural circuits that have overlapping and homogeneous representations should also reflect such a representation at the voxel level. Following this logic, our observations suggest that overlap and homogeneity in neural representation are also unlikely to exist at a more fine-grained scale. However, further investigation will be needed to clarify this point.

## Acknowledgments

We thank Ms. Hitomi Koyama and Ms. Amane Tamai for experimental support. This work was supported by JSPS KAKENHI Grant-in-Aid for Scientific Research B (21H03535 to S.Nishida), JST PRESTO (JPMJPR20C6 to S.Nishida), and NTT Data Japan Corporation (to S.Nishida and S.Nishimoto).

## Extended Data

**Figure 2-1.**
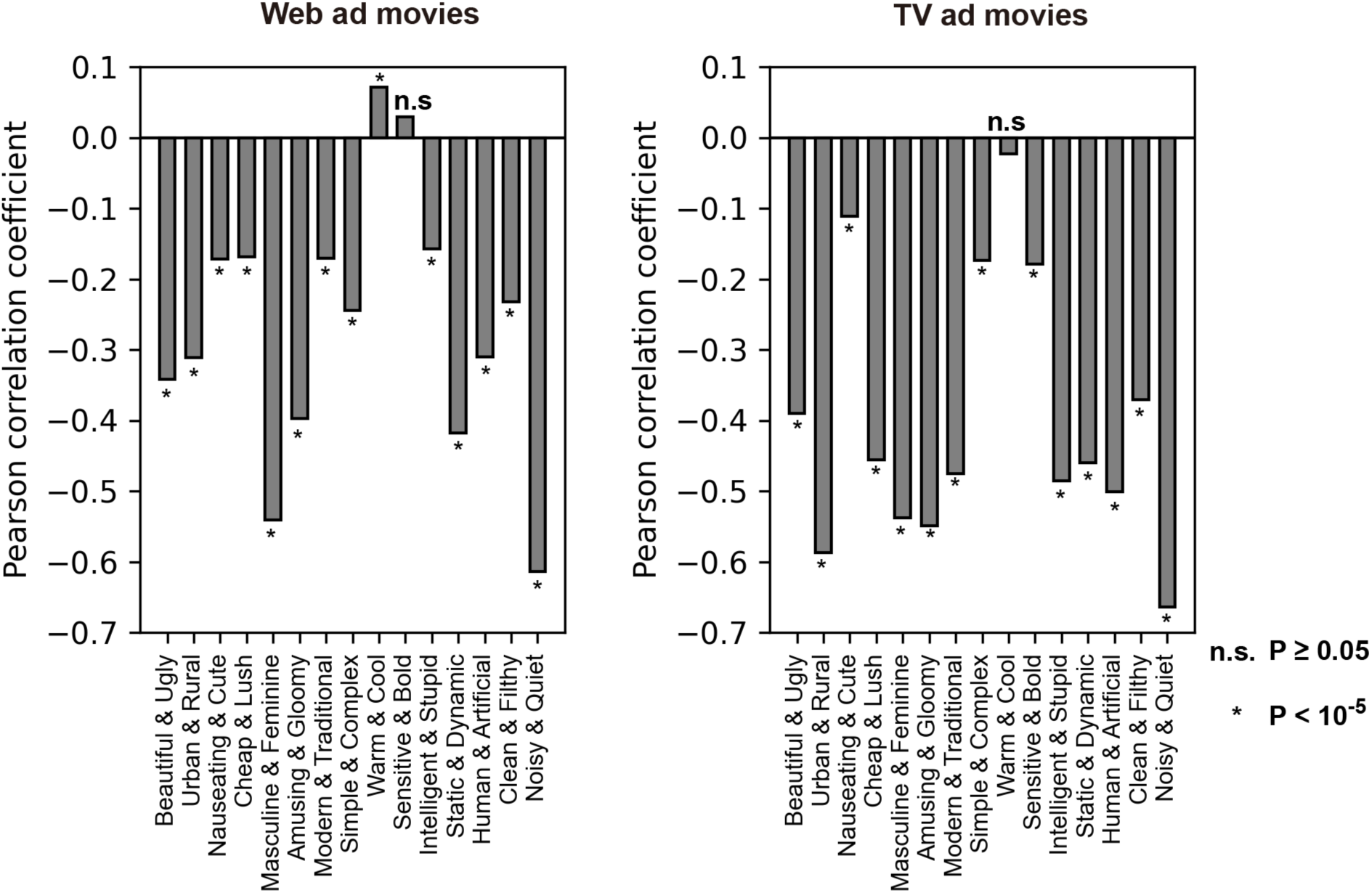
Correlations of ratings for each item pair. Pearson correlation of rating time series for each of the 15 item pairs in each movie set (left, web ad movies; right, TV ad movies). Of all 30 pairs, 27, 1, and 2 pairs showed significantly negative (one-sample t-test, P < 10^−23^, FDR corrected), significantly positive (P < 10^−5^), and non-significant correlations (P ≥ 0.05).

**Figure 3-1.**
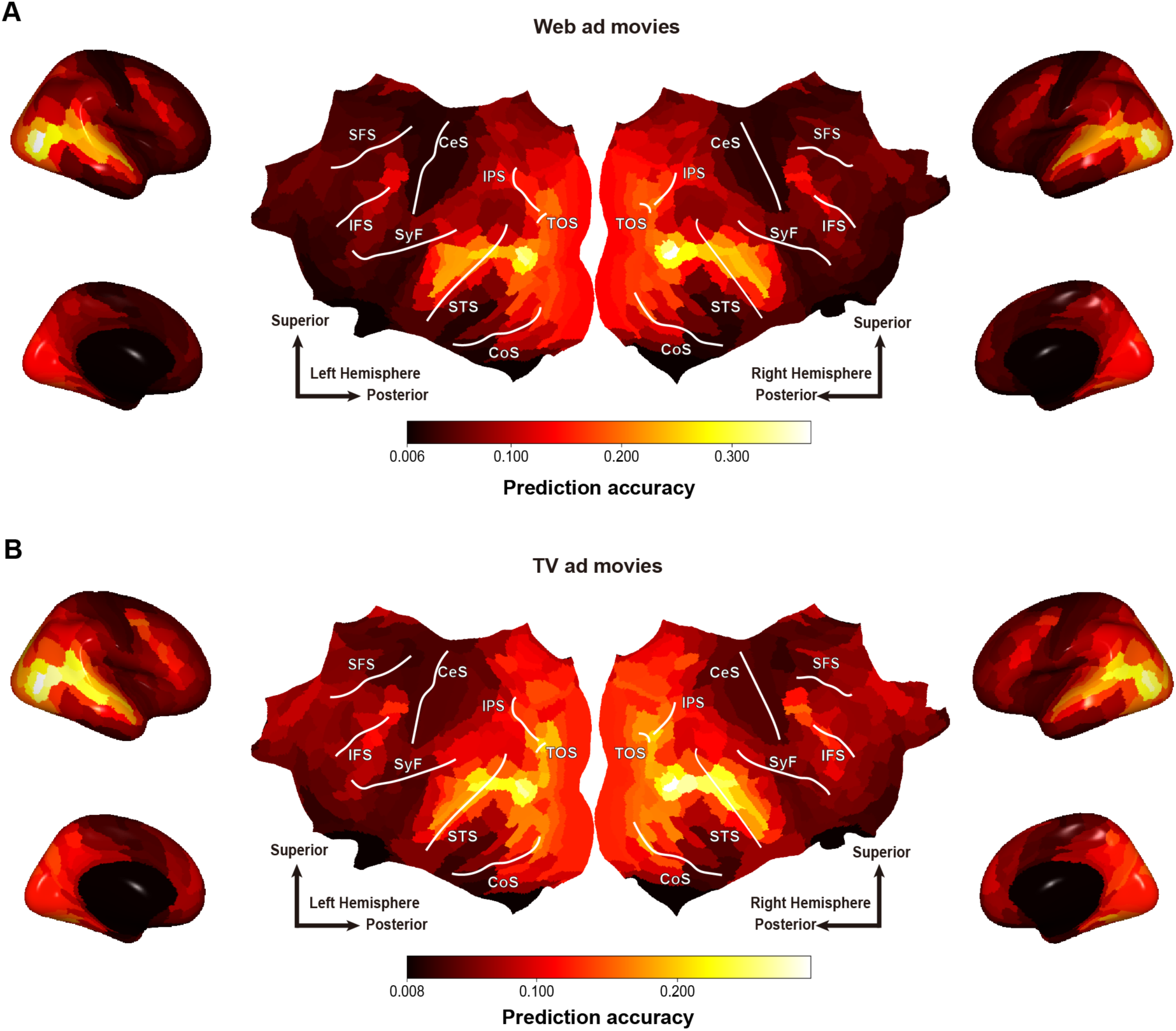
Cortical mapping of prediction accuracy for each movie set. Same as Figure 3B, except that the cortical mapping of prediction accuracy was performed separately for each movie set.

**Figure 3-2.**
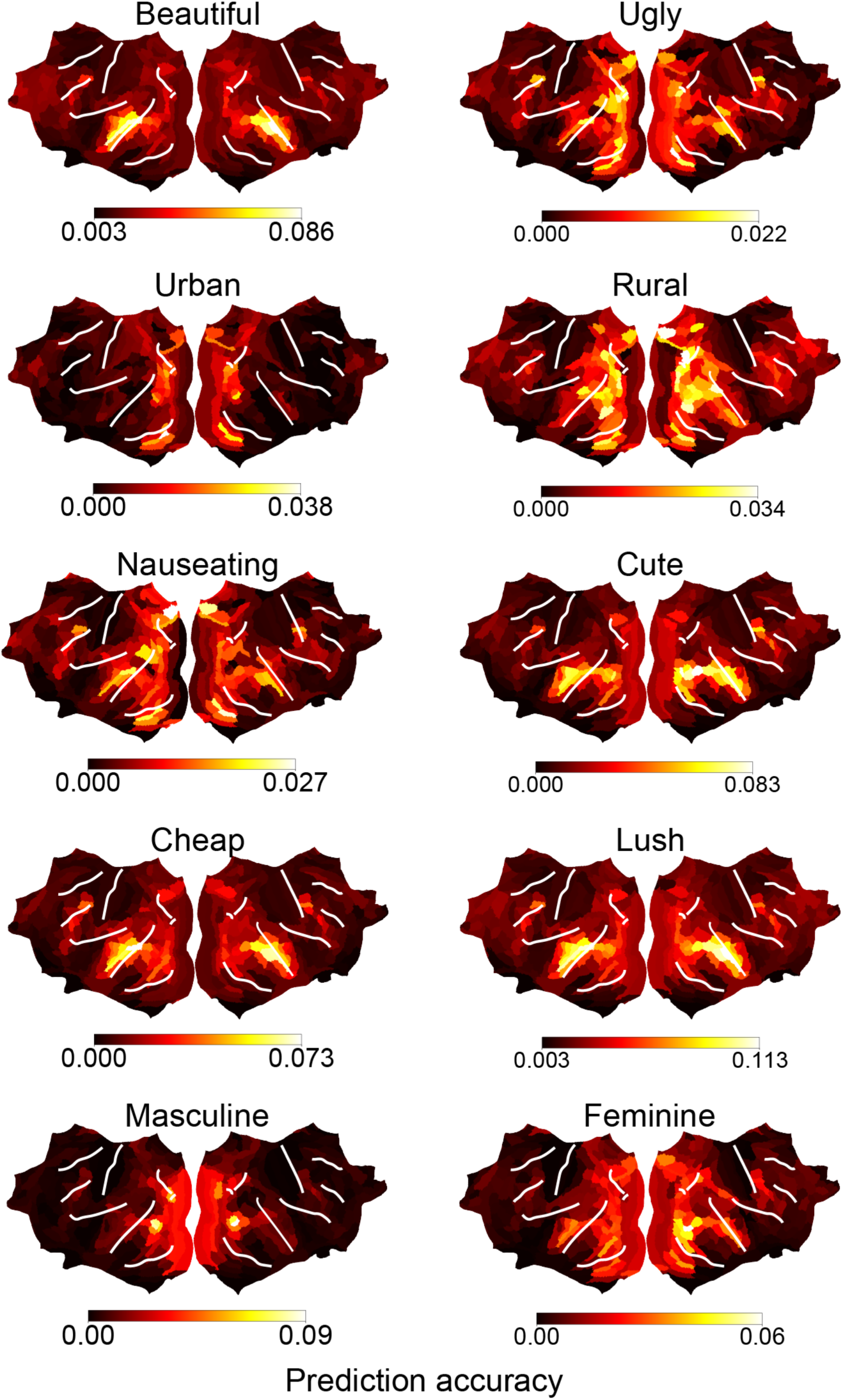
Cortical mapping of prediction accuracy for 10 word items. Region-wise prediction accuracy averaged over participants for each of 10 items mapped onto a cortical surface. The conventions are the same as in Figure 3B.

**Figure 3-3.**
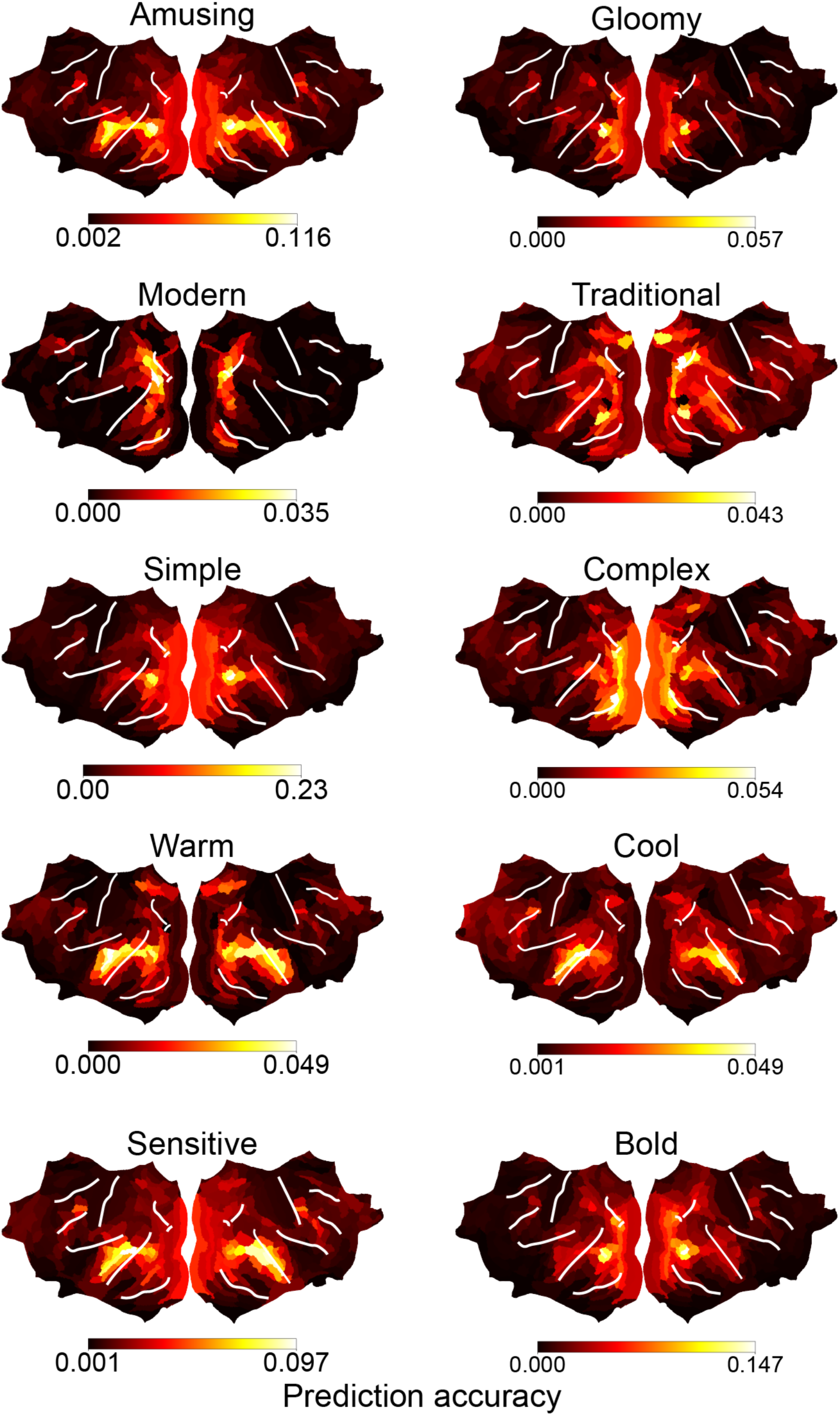
Cortical mapping of prediction accuracy for other 10 word items. Same as Figure 3-2, but other 10 items.

**Figure 3-4.**
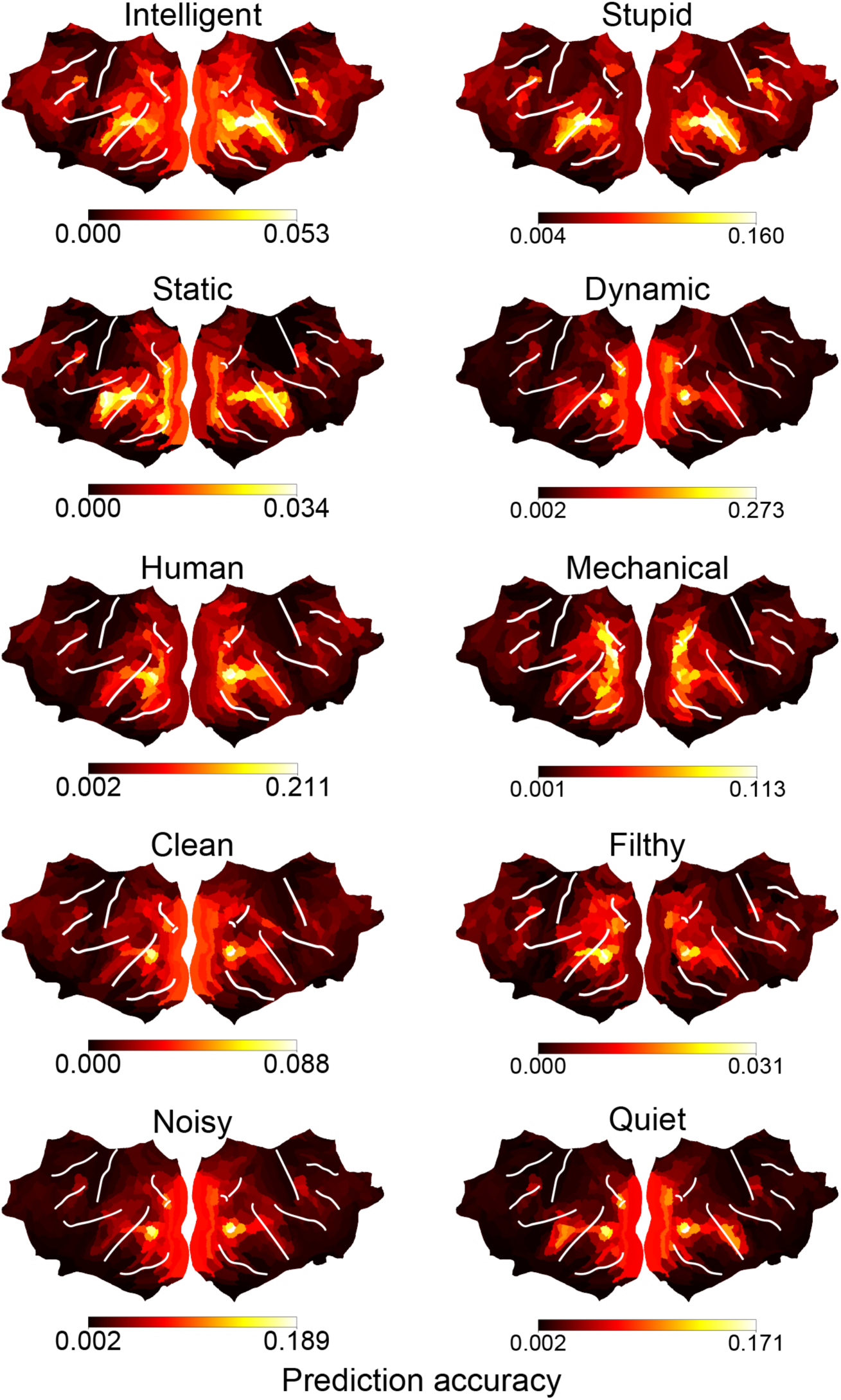
Cortical mapping of prediction accuracy for the other 10 word items. Same as Figure 3-2, but for the other 10 items.

**Figure 5-1.**
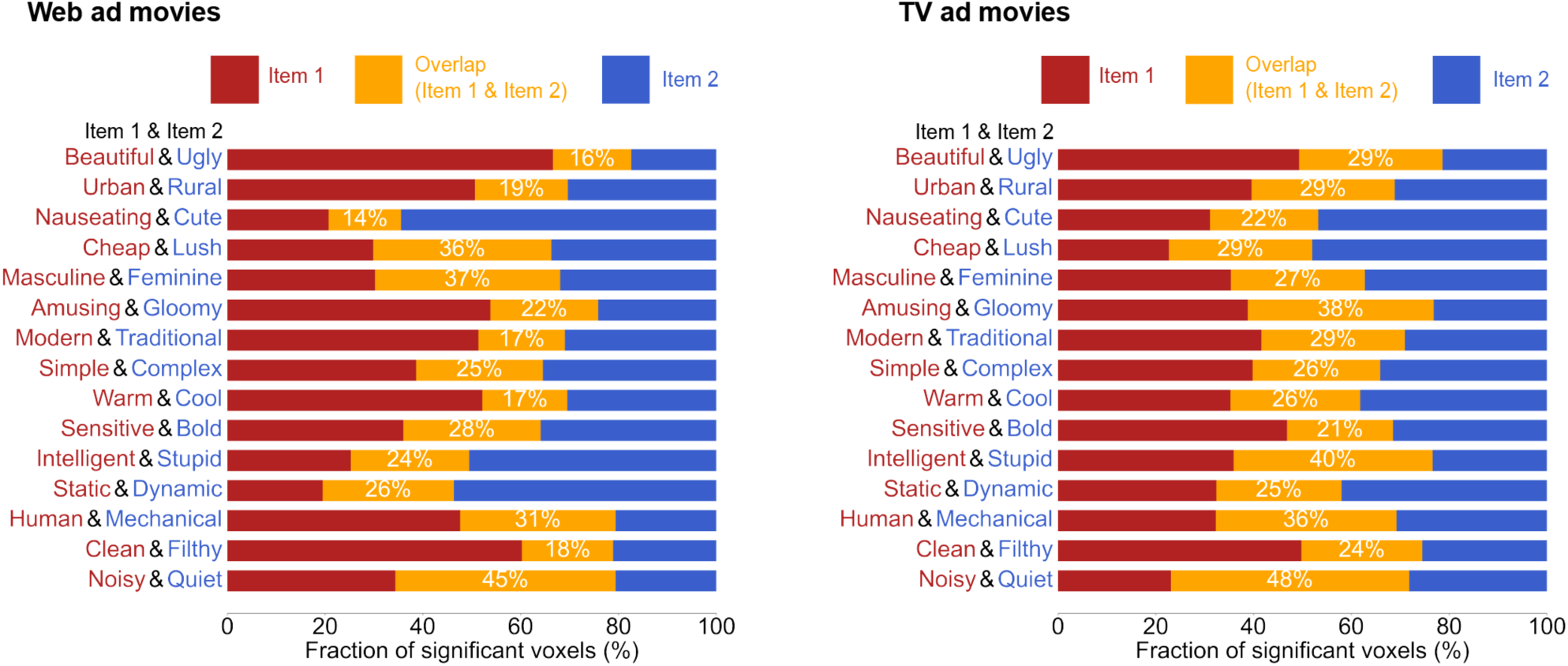
Fraction of different-type predictable voxels for each movie set. Same as Figure 5B, except the fraction was calculated separately for each movie set (left, web ad movies; right, TV ad movies).

**Figure 5-2.**
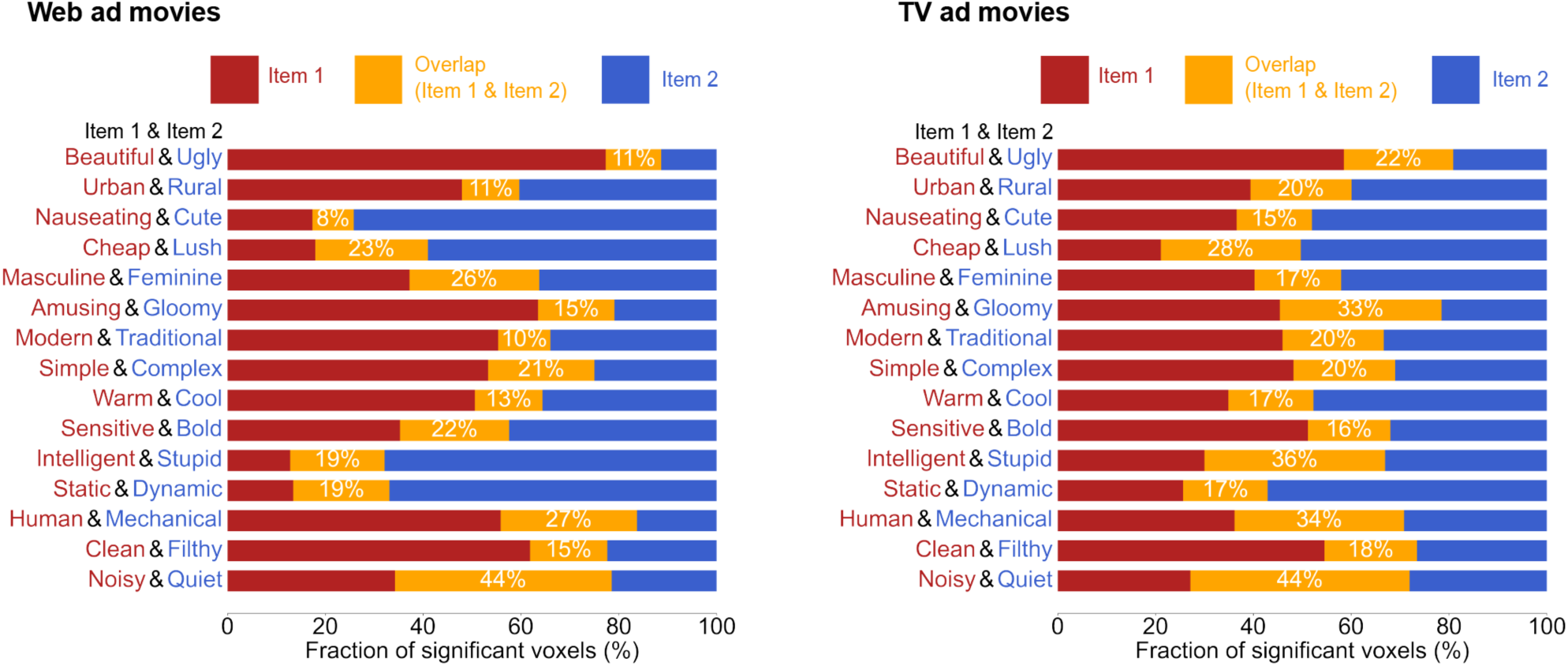
Fraction of different-type predictable voxels for alternative voxelwise modeling. We constructed voxelwise models, each using a five-point scale for one of the 30 items (resulting in 30 models per participant) and calculated the overlap fraction of predictable voxels for each symmetric pair. This fraction, averaged over all participants, is displayed separately for each pair and movie set (left, web ad movies; right, TV ad movies). The conventions are the same as in Figure 5B. Consistent with the original models, the observed fractions were significantly lower than the corresponding chance-level fractions (paired t-test, P < 10^−8^, FDR corrected).

**Figure 6-1.**
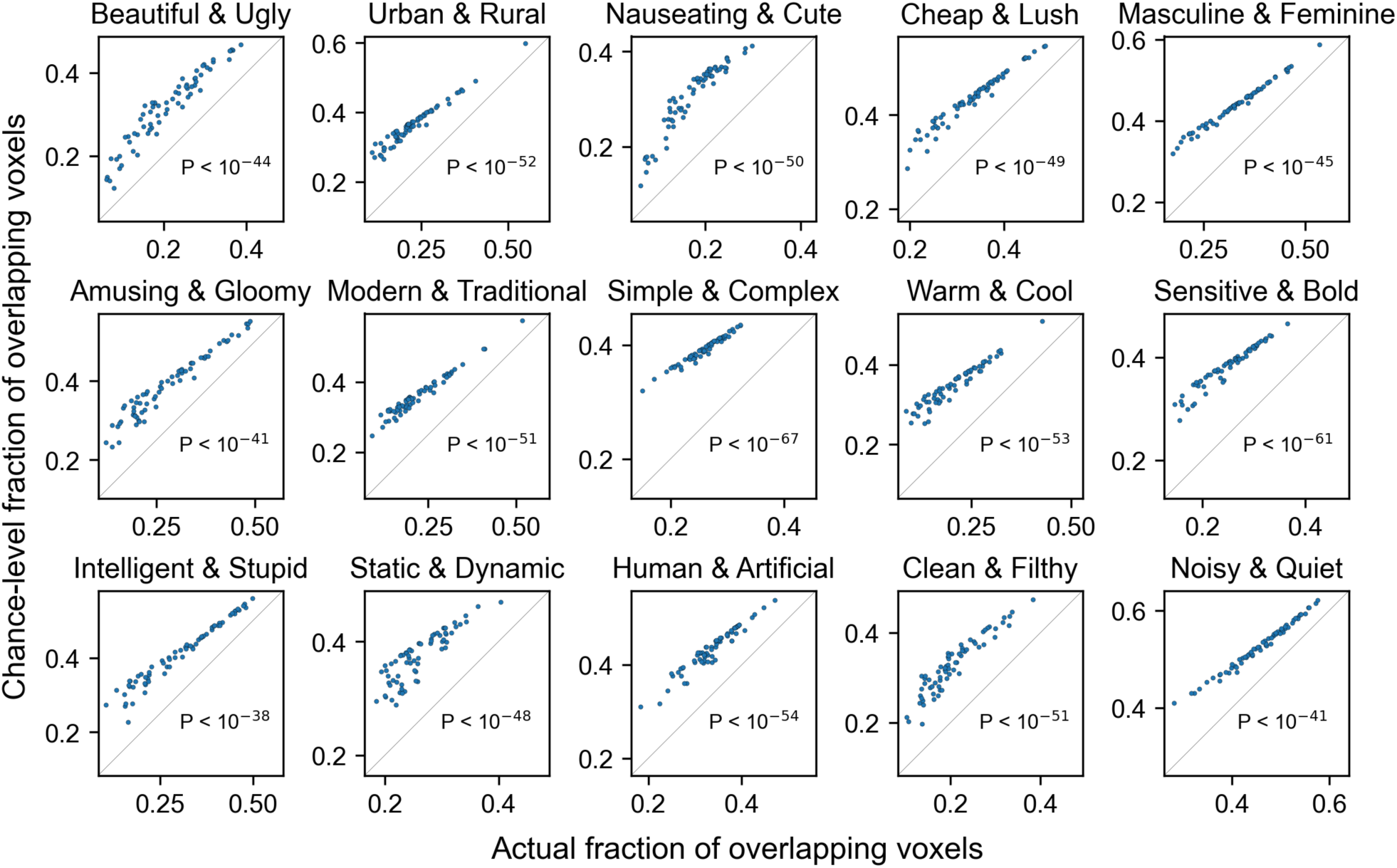
Fraction of overlapping voxels in individual participants and comparison with chance level. Fraction of overlapping voxels obtained from individual participants (x-axis) and the corresponding chance level (y-axis) shown sperately for each item after averaging across movie sets (see Materials and Methods for the calculation of chance level). Each dot in each panel indicates one participant. The fraction for all item pairs was significantly lower than chance level (paired t-test, P < 10^-38^, FDR corrected).

**Figure 7-1.**
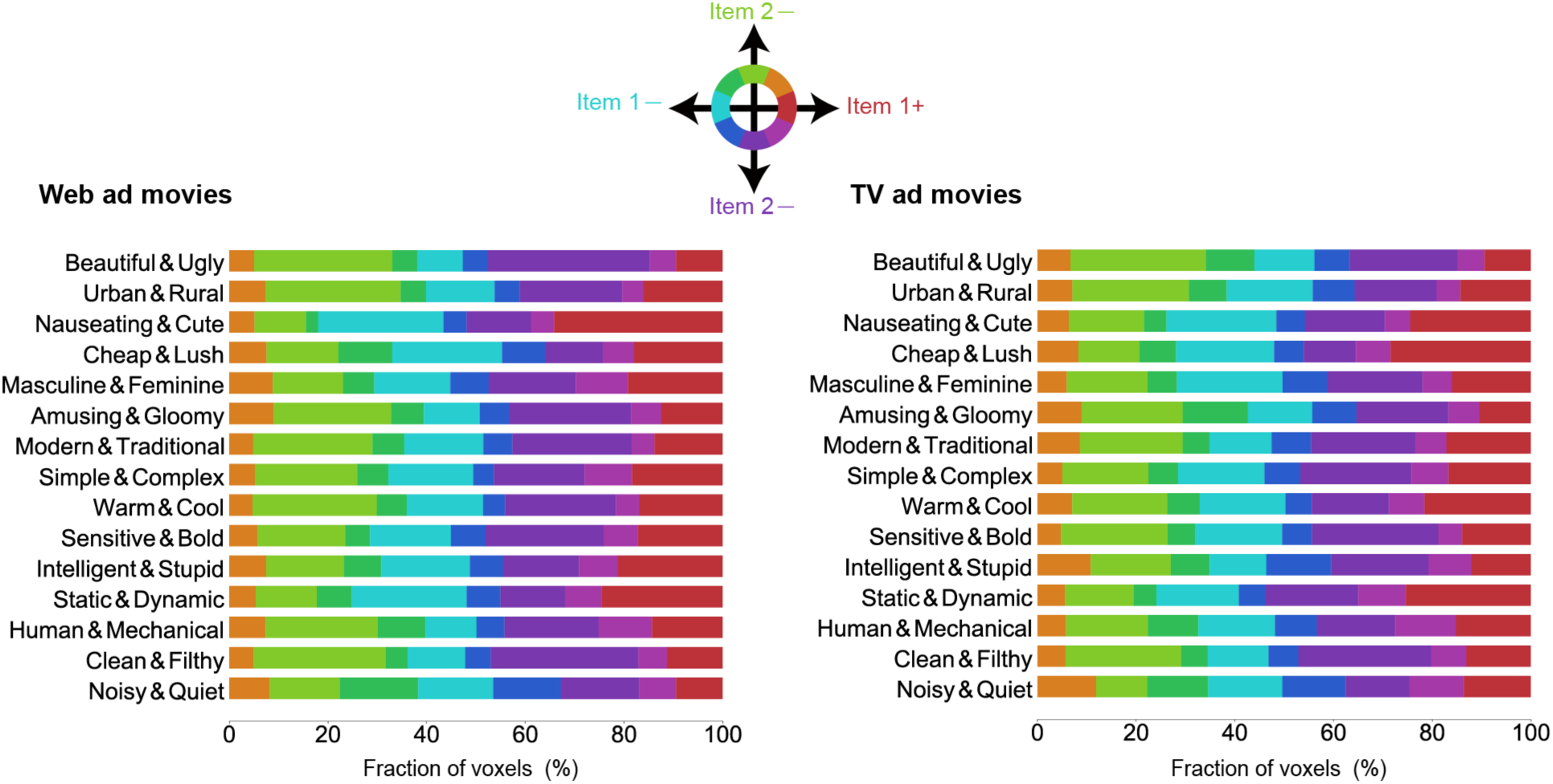
Fraction of different-type selective voxels for each movie set. Same as Figure 7B, except that the fraction was calculated for each movie set (left, web ad movies; right, TV ad movies).

**Figure 7-2.**
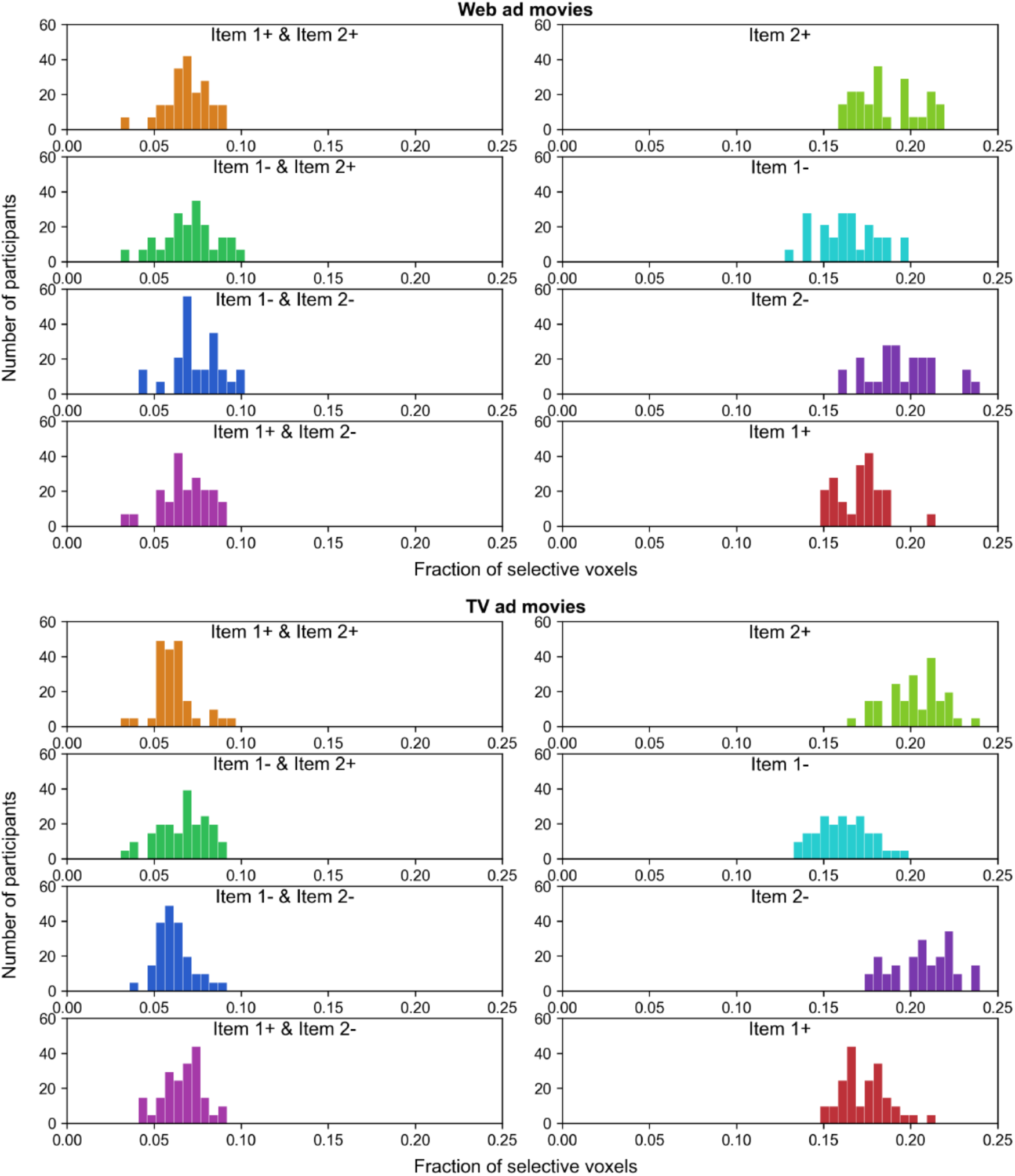
Fraction of different-type selective voxels for individual participants. We classified voxel selectivity into eight types according to model weights (Figure 7). Fraction of selective voxels for individual participants, averaged over item pairs, separately shown for the different types of voxel selectivity (the color in each histogram as in Figure 7) and for different movie sets (top, web ad movies; bottom, TV ad movies).

